# Slow walking synergies reveal a functional role for arm swing asymmetry in healthy adults: a principal component analysis with relation to mechanical work

**DOI:** 10.1101/2020.07.01.182469

**Authors:** David Ó’ Reilly

## Abstract

**Introduction:** The purpose of this study was to reveal a functional role for arm-swing asymmetry during gait in healthy adults. The primary aim was to identify differences in propulsive and collision work between sides at either end of the double-support phase of slow-walking (W_DS_). The secondary aim was to identify differences between sides in propulsive and collision work done at either end of the single-support phase (W_SS_) and the effect of arm-swing asymmetry on this difference. It was hypothesized that differences between sides would be evident during the double-support phase and that these differences would be coherent with differences in single-support control symmetry. It was also hypothesized that left-side dominant arm-swing would reduce the collision work done on the dominant lower-limb side.

**Methods:** A secondary analysis of slow-walking trials of 25 healthy, uninjured adults was undertaken where a principal component analysis of kinematic data was carried out to generate the movement synergies (PM_k_). Independent variables included the tightness of neuromuscular control (N_k_) which was formulated from the first PM_k_ and arm-swing asymmetry which was quantified using the directional Arm-swing asymmetry index (dASI). Dependent variables included the difference between double-support collision and propulsive work (W_DS_) and a ratio consisting of the difference between single-support collision and propulsive work of both sides (W_SS_). A linear mixed-effects model was utilized for aim 1 while a multiple linear regression analysis was undertaken for aim 2.

**Results:** Healthy adult gait was accompanied by a left-side dominant arm-swing on average as seen elsewhere. For aim 1, N_k_ demonstrated a significant negative effect on W_DS_ while sidedness had a direct negative effect and indirect positive effect through N_k_ on W_DS_. The most notable finding was the effect of a crossover interaction between dASI and N_k_ which demonstrated a highly significant positive effect on W_ss_. All main-effects in aim 2 were in the hypothesized direction but were insignificant.

**Interpretation:** The aim 1 hypothesis was supported while the aim 2 hypothesis was not supported. N_k_ exhibited opposing signs between ipsilateral and contralateral WBAM regulation, revealing a differential control strategy while the effect of sidedness on W_DS_ was evident. The findings from aim 2 describe a relationship between arm-swing asymmetry and the magnitude of lower-limb mechanical work asymmetry that is cohesive with the sidedness effect found in aim 1. Individuals with left-side dominant arm-swing had an increased collision work indicative of a lateralised preference for WBAM regulation. Evidence was therefore put forward that arm-swing asymmetry during gait is related to footedness. Future studies should look to formally confirm this finding. Implications for further research into dynamic balance control mechanisms are also discussed.

**Highlights:**

- Left-side dominant arm-swing was found to be related to the degree of lower-limb mechanical work asymmetry.
- The relationship between arm-swing asymmetry and lower-limb mechanical work symmetry was explained by a moderating effect of neuromuscular control.
- A differential control on single-and double-support phases was demonstrated by the neuromuscular system, supporting previous studies and this control may be heavily influenced by sidedness.

## 1. Introduction

Walking is an activity central to an individuals’ everyday life and functional independence. As such, rehabilitative and movement science research dedicates a significant proportion of its focus on this activity as acquired knowledge can inform important intervention design and surgical decision-making (1, 2). Gait is said to be analogous to an inverted-pendulum like motion where one limb supports the bodyweight and the other moves towards and past this supporting limb to advance the whole body (3). The upper-limbs also simultaneously swing in asynchrony to this lower-limb motion. Evidence suggests that arm-swing is required to minimize energy expenditure and optimise dynamic stability (4, 5). Energy efficiency can be improved for example through elevation of the arm at terminal-swing, where the trunk is lifted upwards allowing for reduced collision work at heel-strike contralaterally (6). Dynamic stability can be optimised by medio-lateral extension when a perturbation is experienced or the guard posture in anticipation of a fall for example (7, 8).

Arm-swing symmetry is relevant in assessing the early presentation of Parkinson’s disease and deficits in gait stability as notable asymmetries are often present (7, 9, 10). The presence of arm-swing asymmetry within healthy populations has been noted in the literature (10), suggesting that the aforementioned clinical evaluations could potentially be confounded by pre-existing asymmetries. The aetiology for arm-swing symmetry in healthy populations is not well understood. In patient populations, it has been related to inherent asymmetries in the control of other body segments (e.g. hemiparetic gait) (7). This upper-limb asymmetry counteracts increased angular momentum within the lower-limbs (11, 12), enabling a more stable gait pattern. Strong studies have purported this asymmetry to be linked to handedness in Parkinson’s disease (13, 14). Studies investigating the relationship between arm-swing asymmetry and handedness among healthy adults however found no association despite the majority of healthy adults demonstrating a left-side dominant arm-swing (15, 16). Nonetheless when a left-lateralised task was added in a dual-task walking condition, noticeable increases in arm-asymmetry were found indicating the role of cerebral lateralisation (10, 15).

It is of frequent practice in research to challenge the central nervous system (CNS) of participants for example by creating dual-task conditions (15), inducing a perturbation (17), standing on unstable support surfaces (18) or walking more slowly/quickly than usual (19). Challenging subjects to intentionally walk slower than normal induces lower inter-limb coordination and increased attentional requirements (20). Through such challenging methodologies, useful insight can be gained into the mechanisms underlying human movement. Coordinated movement is highly complex and redundant in that the CNS has more ways than needed for carrying out a given task. This redundancy was originally perceived as the ‘problem of redundancy’ as it was thought the CNS constrained the multiple degrees-of-freedom into a task relevant movement (21). More recently, this problem has been re-interpreted in a more positive light as motor abundancy where the CNS selects a near-optimal movement strategy from the range of possible strategies and intervenes thereafter only when the movement goes outside a task-relevant range (22, 23). Efficient selection of movement strategies is carried out via a modular control of muscle activations, known as muscle synergies at the level of the spine within central pattern generators (CPGs) (24).

Robert et al. (2009) through the uncontrolled manifold approach, revealed a modulated control of whole-body angular momentum (WBAM) during gait that was noticeably different during the double-and single-support phases. The double-support phase was characterised as a closed-loop system where postural corrections could take place. The single-support phase was dedicated towards providing step-to-step reproducibility of the WBAM. These opposing roles for WBAM regulation indicate that the neuromuscular system actively intervenes during the double-support phase while only intervening when necessary during the single-support phase. This active intervention is necessary to allow for smooth, coordinated step-to-step transitions while also preventing the accumulation of small perturbations (3, 26, 27). The majority of gait energy expenditure is said to be caused by the mechanical work involved in step-to-step transitions (i.e. propulsion and collision work) (28). These opposing forces interact to stabilise the trajectory of the net vertical force produced during step-to-step transitions and are central to the internal work conducted during gait (26). Using passive gait models, arm swing has demonstrated a role in controlling the magnitude of vertical force thus indicating a pivotal role for opposing upper-limb motion (29). To date it has not been investigated to what extent the mechanical work during the step-to-step transitions of gait is symmetrical across sides in healthy adults and also to what extent arm-swing asymmetry plays a role in this.

Specific synergy extraction protocols for motion capture data have been formulated that involve the use of principal component analysis (PCA) (30). PCA is a machine-learning algorithm which effectively reduces high-dimensional data down to a smaller number of orthogonal vector components known as ‘*Principal movements’* (PM_k_) (30). Each PMk can be interpreted as correlated marker movements or synergies with higher-order PM_k_ explaining less variance and representing more subtle movements. Successful uses for this PCA protocol thus far include identifying differences between healthy and knee osteoarthritic gait (31), quantifying sporting techniques (32) and determining age and leg dominance effects on postural control (33, 34). The described PM_k_ can be projected onto a posture space, and in doing so can be represented as ‘*Principal positions’* with respect to time (PP_k_). Similar to Newton’s laws of motion, these PP_k_ can be differentiated into their 1^st^- and 2^nd^-order derivatives, ‘*Principal velocities’* (PV_k_) and ‘*Principal accelerations’* (PA_k_) (30). The PA_k_have been of particular interest as they are thought to represent the action of the neuromuscular system (34, 35). Among other potential variables, the tightness of neuromuscular control on PM_k_ (N_k_) can be determined by the number of zero-crossings (changes in direction) in the corresponding PA_k_ (34, 36).

The purpose of the current study is to investigate the mechanical work conducted during step-to-step transitions of very slow walking in healthy adults contralaterally (double-support) and ipsilaterally (single-support) while considering the effect of sidedness and arm-swing asymmetry respectively also. Through this analysis, an understanding of the mechanical work regulating WBAM and how it is affected by sidedness and arm-swing asymmetry can be gained. The following aims will be undertaken to fulfil this purpose: 1) determine the relationship between N_k_ and differences in collision and propulsive work conducted contralaterally in the context of sidedness. 2) establish the relationship between N_k_ and arm-swing asymmetry with collision and propulsive work conducted ipsilaterally during gait. From this, a functional role for arm-swing asymmetry can be revealed and whether sidedness may play a role. It is hypothesised for aim 1 that N_k_ will have a negative association in that greater tightness of control will reduce the magnitude of collision work necessary for WBAM regulation however this effect will also interact significantly with limb-side. Conversely for aim 2, it is hypothesized that N_k_ will have a positive effect on ipsilateral collision work in that greater tightness of neuromuscular control will predict greater ipsilateral collision work. It is also hypothesized that arm-swing asymmetry will regulate WBAM by reducing collision work in a manner that is cohesive with the sidedness effect found in aim 1.

## 2. Methods

### 2.1 Secondary data analysis

Three-dimensional marker trajectories from slow-walking trials of 25 healthy, injury-free adults were taken from a publicly available open-source dataset published in *Nature* (37). This motion capture data was generated using a 10-camera optoelectronic system sampled at 100 Hz (OQUS4, Qualisys, Sweden) where a 52 markers biomechanical model setup was utilized and marker trajectories were filtered with a 4^th^-order Butterworth low-pass filter at cut-off frequency of 6Hz (38). In the current study, this marker setup was simplified to 36 markers. 13 male and 12 female participants (Age: 32.88±10.6, Height: 1.72±0.1, Weight: 71.4±11.2, BMI: 24.04±2.4) were asked to walk at a speed between 0m/s – 0.4m/s (corresponding to a ‘household ambulator’ (39)) that was coordinated by a metronome on a 10-metre straight and level walkway. Participants were selected for the current study if they had at least 4 out of the possible 5 trials at this walking speed while inclusion criteria in the source study meant only individuals with a leg length difference of less than 1.5% of their height (approx. 0.03m). One right and left gait-cycle were captured in each trial. This data was collected originally using Qualisys Track Manager software (QTM 2.8.1065, Qualisys, Sweden). For further details on the data collection procedure, please see (37).

### 2.2 Synergy extraction

All of the beforementioned procedures were carried out using PManalyzer (40), a Matlab graphical user interface specifically designed for PM_k_ computation and visualization (Matlab (R2019B), Natick, Massachusetts: The MathWorks Inc.). XYZ marker coordinate trajectories from each trial were concatenated and the length of these time-series were normalised to a median range value of 2220 data points for each participant. The individual matrices were then pooled into one 55,500 x 108 input matrix (100 Hz [Sampling rate] x 2,220 data points [Trial duration] x 25 [Number of participants] x 108 [Marker coordinates]) to allow for direct comparisons between participants. In order to eliminate anthropometric differences across participants, this input matrix was firstly transposed so that each time-frame represents a posture vector which were then centred by subtracting these vectors from their respective averages, creating postural deviation vectors. These postural deviation vectors were also centred towards the centre-of-mass to avoid the inclusion of body displacements within the PM_k_ (32). Finally, the postural deviation vectors were normalised by their mean Euclidean distances to ensure an equal contribution of all participants to the PCA output. A PCA algorithm via singular value decomposition then converted this input matrix into a covariance matrix of PM _k_ (41).

### 2.3 Independent variable computation

From this protocol, the PM_k_ were projected onto the mean posture space and PP_k_ were derived. These PP_k_ were inspected using a Fourier analysis for noise. Further filtration with a 3^rd^-order Butterworth low-pass filter at a cut-off of 10Hz was deemed necessary to prevent the amplification of noise with differentiation. These filtered PP_k_ were then differentiated to their 2_nd_-order derivatives PA_k_. A leave-one out cross-validation revealed that the first 4 PM_k_ did not change by >15° and these were deemed valid (34). The first PM explaining 61.7% of the overall variance was chosen for further analysis only as this primary PM has been identified in previous studies as the basic inverted-pendulum motion of gait and as such is of specific relevance to this study’s aims (17, 42). With this primary PM the number of zero-crossings in its corresponding PA_k_ time-series were counted for each participant (34, 36), thus formulating the independent variable N_k_ for further analysis.

The directional Arm-swing asymmetry index (dASI) was formulated to determine the degree of arm-swing asymmetry during gait. The medial wrist marker trajectories for both sides in the sagittal plane were extracted and the range of motion (ROM) of these trajectories with respect to the participants centre-of-mass were taken (16). The centre-of-mass in this case was specified as the centroid of a geometric triangle made up of the two anterior superior iliac-spine markers and the midpoint of the two posterior superior iliac-spine markers (43). Equation 1 below illustrates how dASI was calculated using these arm-swing ROMs where ***L*** is the ROM of the left-arm and ***R*** the right-side (10, 15). A positive dASI value indicates left-side dominant arm-swing while vice versa.

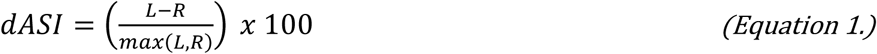

### 2.3 Dependent variable computation

Using the CusToM toolbox (Customizable Toolbox for Musculoskeletal simulation) in Matlab (44), a biomechanical model consisting of 17 rigid body segments linked by 14 joints was created. This model was geometrically scaled to each participants height and weight while segment masses and inertia were calibrated using Body Segment Inertia Parameters (45). Using the Levenberg-Marquardt algorithm, whole-body 3D inverse kinematics were extracted followed by joint torques. Heel-strike and toe-off events were detecting using a kinematic method (46) and the datapoints for these events were taken from the kinematics and joint torque time-series for further analysis. Kinematic time-series were then converted to angular displacement vectors. The average joint work done 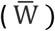 across trials at both gait events on each side was calculated as the average joint angular displacement 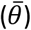 multiplied by the corresponding average joint torque 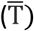 (Equation 2). The corresponding hip- and ankle-joint data for heel-strike and toe-off events respectively were then extracted. These events data would now represent collision and propulsive work respectively.

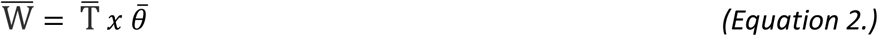

The following variables were then created to capture the mechanical work involved in step-to-step transitions both contralaterally and ipsilaterally. For aim 1, the difference between 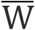 for heel-strike event 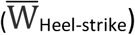 and 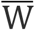 for contralateral toe-off events 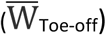 was calculated on both sides (Equation 3). This dependent variable W_DS_ allows one to determine the typical differences in propulsive work at toe-off and collision work done contralaterally at heel-strike, providing a representation of double-support phase mechanical work. For aim 2, the absolute difference between right-side 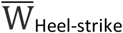 and right-side 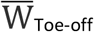 was divided by the equivalent absolute difference on the left-side (Equation 4). This ratio W_SS_compares the symmetry of propulsive and collision work done ipsilaterally between sides, providing a representation of single-support phase mechanical work and how they compare across sides. A higher W_ss_ value indicates a preference for right-side collision work. Contrasting the magnitude of collision work between sides during very slow walking will be descriptive of the healthy participants symmetry in lower-limb motor control. Figure 1 below further illustrates W_DS_ and W_ss_ where B denotes 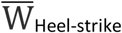 and P signifies 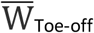.

**Figure 1:**
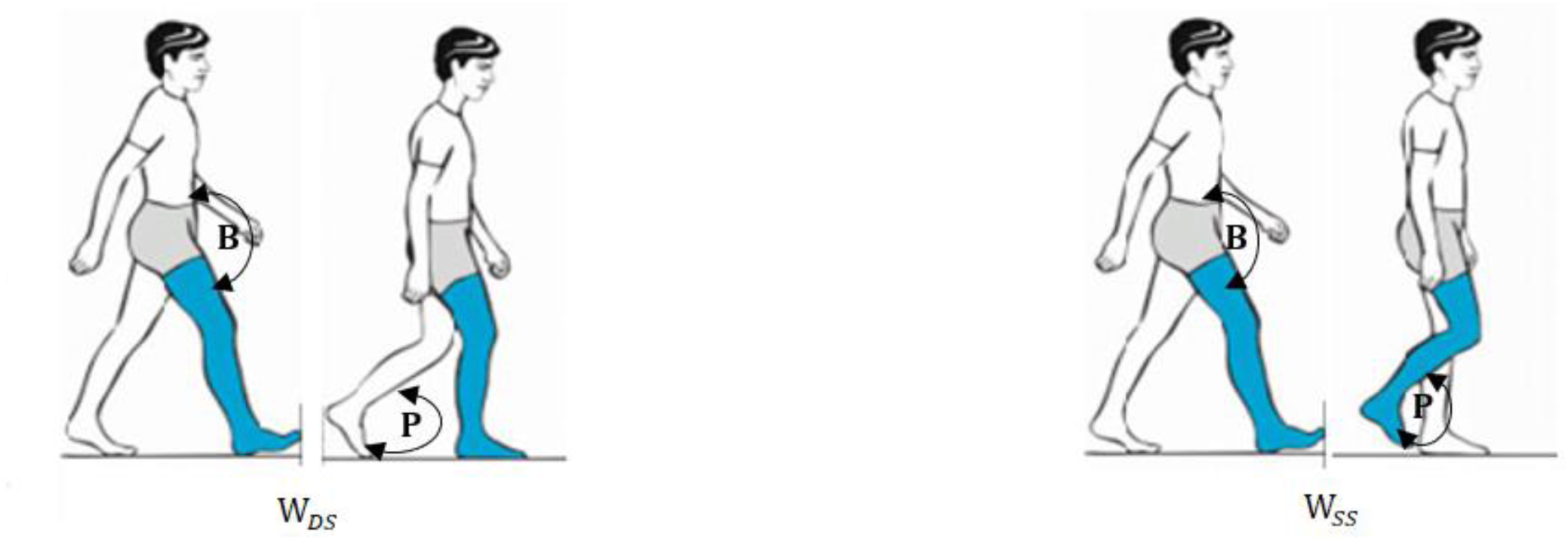
An illustration of the mechanical work being compared contralaterally (W_DS_) and ipsilaterally (W_SS_) during step-to-step transitions. **B** denotes the braking work conducted by the hip joint at heel-strike (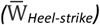) while **P** represents the propulsive work done by the ankle joint at toe-off 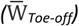.

**Figure 2:**
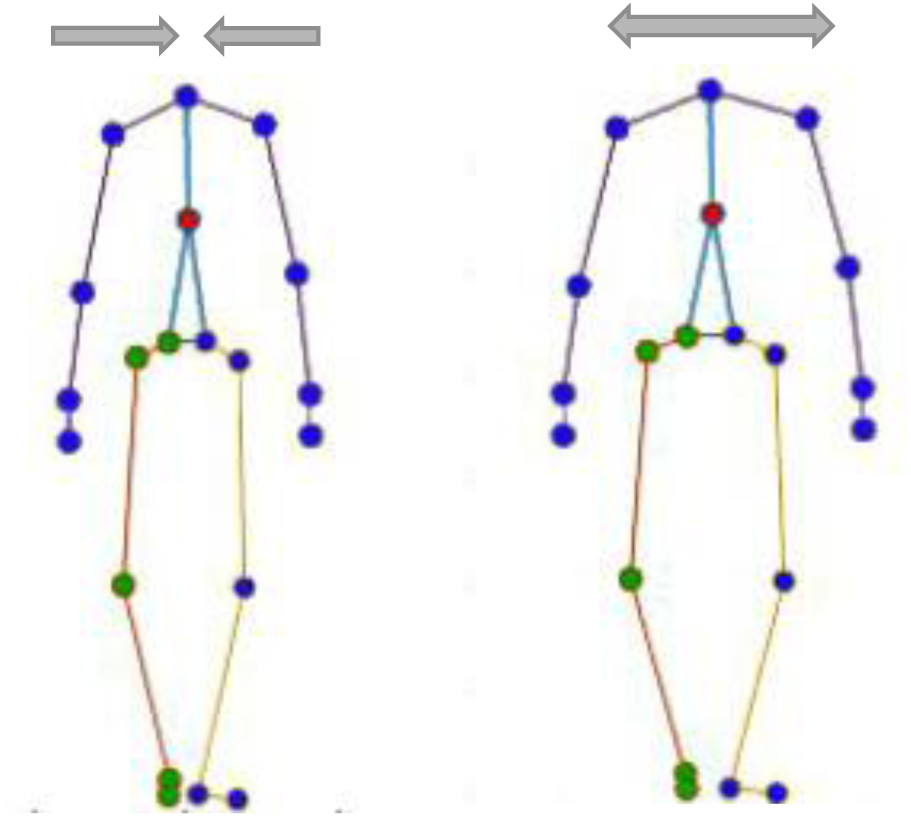
Frontal plane graphical representations of the primary PM.

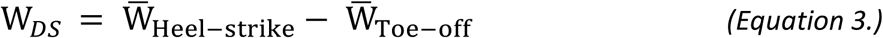

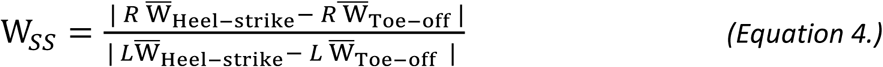

### 2.4 Statistical analysis

As W_DS_ determined differences on both sides and these differences were not incorporated into the one observation as is the case with W_SS_, a linear mixed-effects model was utilised for aim 1 to determine the effect of groupings within the data. Equation 5 below illustrates the formula for this analysis in R syntax where (*N*_*k*_ |Participant) is a random-intercepts and slopes term allowing both the start point and slope of the regression line to vary between-participants. *N*_*k*_, *Side* and *N*_*k*_: *Side* are fixed-effects terms, similar to ordinary-least squares regression. *N*_*k*_: *Side* represents the interaction between *N*_*k*_ and the side of W_DS_. Mixed-effects modelling was carried out using the lme4 package in R (47). Side was coded as Right=1 and Left=2 so a negative effect in this instance would represent right-side dominance.

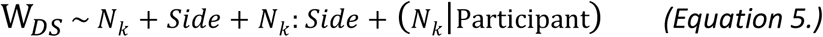

For aim 2, a multiple linear-regression analysis was undertaken in SPSS (IBM SPSS Statistics, Version 26. Armonk, NY: IBM Corp) in which *N*_*k*_, *dASI* and the interaction term (N_k_ x *dASI*) for these two variables were modelled against W_SS_. Significance was set a priori at p<0.05 for both aims.

## 3. Results

### 3.1 Slow-walking synergies

As the walking-speed of healthy adults in this analysis was very slow (0.1m/s – 0.4m/s), the primary PM_k_ used in this study captured the predominant movement at this very slow walking-speed, that of upper-and lower-limb medio-lateral motion. This insight was gained by reversing the PM_k_ normalistion procedure and projecting them onto the posture space which can then be graphically represented. Figure 1 exemplifies this interpretation in the frontral plane with the upper- and lower-limbs moving synchronously towards and away from the bodys center.

Table 1 provides a descriptive outline of the various variables of interest in the current study. Participants were typically left-side dominant in terms of arm-swing (dASI: 11.06±28.86) but ranged widely from -49.66 to +51.32 for this metric. The number of direction changes in the primary PA component (number of ‘zero-crossings’) was typically 109.76±18.15 and ranged widely also from 77 – 161. W_ss_ varied widely from -308.49 - +564.07 indicating a significant between-participant variability while W_DS_ varied to a much lower extent (−9.16 – +11.80).

**Table 1:** Descriptive statistics for the variables of interest.

| Variable | Mean (SD) | Minimum | Maximum |
| --- | --- | --- | --- |
| $dASI$ | $11.06 \pm 28.86$ | -49.66 | 51.32 |
| $N_k$ | $109.76 \pm 18.15$ | 77 | 161 |
| $W_{SS}$ | $11.52 \pm 129.56$ | -308.49 | 564.07 |
| $W_{DS}$ | $0.77 \pm 3.77$ | -9.16 | 11.80 |

### 1.1 Aim 1

Table 2 describes the output from a linear mixed-effect model described in Equation 5 of the methods section. *N*_*k*_ demonstrated a significant negative effect on W_DS_ (β= -57.54±20.87, df= 38.73, p<0.01) while sidedness also demonstrated a large negative effect with a high degree of variability (β= -2973.98±1425.11, df=32.79, p<0.05). Conversely the interaction term Equation 5 of the methods section: *Side* demonstrated a significant positive on W_DS_ (β= 29.6±12.81, df=32.79, p<0.05). Together, these results indicate that with greater tightness of neuromuscular control, W_DS_ is reduced so that negative work is reduced relative to propulsive work. The right-side double-support phase appears to be favoured in terms of WBAM regulation in this cohort as a greater degree of collision work was done on this side. Moreover when *N*_*k*_ is high on the left-side, the effect of sidedness is reduced as collision work increases relative to propulsive work during left-side double-support, thus sharing the role of WBAM regulation more evenly across sides. The within-participant clustering for intercepts was large with a standard deviation (SD) of 101.42 while random slopes clustered within-subject with a SD= 3.59 indicating the effect of *N*_*k*_ may differ to a degree between sides as was evidenced in the contrasting effect signs. Between-participant variance was highest in this sample at SD=821.96 indicating that the sample overall behaved heterogeneously during slow-walking trials in terms of *N*_*k*_.

**Table 2:**
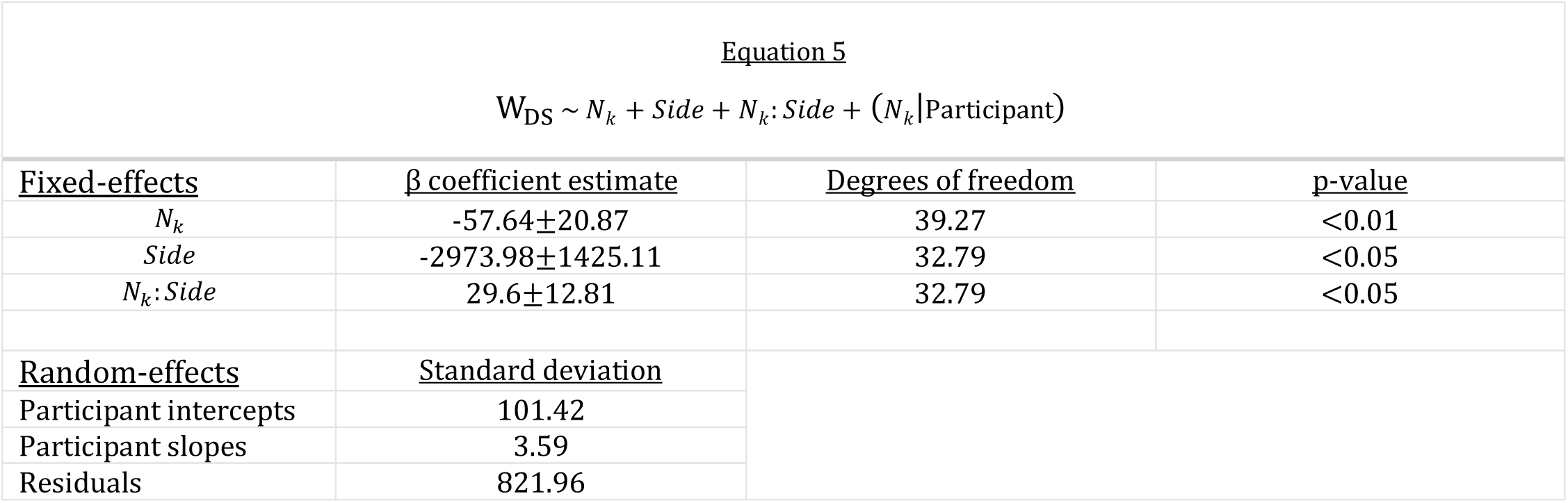
Findings from a linear mixed effects regression analysis with Equation 5 as the input.

### 1.2 Aim 2

A multiple-linear regression analysis of *N*_*k*_, dASI and their interaction term (*N*_*k*_ x dASI) revealed insignificant direct effects for both *N*_*k*_ (β =0.038±0.25, p>0.2) and dASI (β= -0.037±0.162, p>0.3) but a highly significant indirect effect for *N*_*k*_ x dASI (β= 0.96±0.00) was found. Table 3 illustrates these findings further with an R-squared value of 0.977 overall for this model. From this, it can be understood that with greater tightness of neuromuscular control in the presence of a left-side dominant arm-swing, WBAM regulation is made significantly more asymmetrical. In this cohort, the asymmetry favoured the right-side. Moreover, as both direct effects were insignificant the interaction term can be interpreted as a cross-over interaction in that when N_k_, dASI or both are relatively low, the effect of this interaction is the opposite of when both variables are high.

**Table 3:**
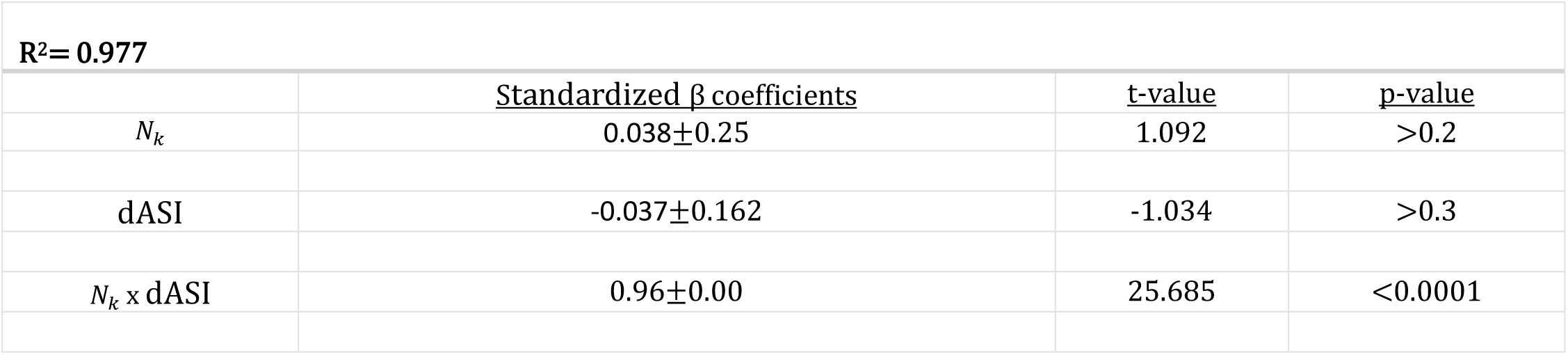
Findings from a multiple-linear regression analysis with *W*_*ss*_ as the dependent variable.

| $R^2 = 0.977$ | | | |
| --- | --- | --- | --- |
| | Standardized $\beta$ coefficients | t-value | p-value |
| $N_k$ | $0.038 \pm 0.25$ | 1.092 | >0.2 |
| dASI | $-0.037 \pm 0.162$ | -1.034 | >0.3 |
| $N_k \times dASI$ | $0.96 \pm 0.00$ | 25.685 | <0.0001 |

## 2. Discussion

The purpose of this study was to reveal a functional role for arm-swing asymmetry in healthy adults via slow-walking synergies. This was conducted by firstly investigating the relationship between the tightness of neuromuscular control and mechanical work conducted contralaterally (double-support phase) in the context of sidedness. It was hypothesized that a tendency for mechanical work to be dictated predominantly by one side would be present while higher N_k_ would be beneficial for mechanical work efficiency during this phase. In aim 2, the association between arms-wing asymmetry and tightness of neuromuscular with ipsilateral mechanical work (single-support phase) was investigated. It was also hypothesized that left-side dominant arm-swing would reduce right-side collision work and therefore be beneficial to mechanical work efficiency while this effect would be cohesive with the sidedness effect found in aim 1. The first PM was taken and the number of ‘zero-crossings’ (N_k_) in the corresponding PA component for each participant was extracted. The findings of this study were in agreement with the aim 1 hypothesis but not so in the case of aim 2 hypothesis. In aim 1, N_k_ demonstrated a negative effect on W_DS_ while side was also significantly negative in its effect with participants favouring right-side WBAM regulation during double-support and N_k_ was beneficial in reducing the degree of collision work conducted. For the secondary aim, the direct effects of N_k_ and dASI were insignificant in their effect on W_ss_. However, their interaction revealed a highly significant positive effect demonstrating that with a high level of N_k_ increased the relationship between dASI and lower-limb mechanical work asymmetry (higher left-side dominant arm-swing related to right-side dominant mechanical work).

### 2.1 A functional role of arm-swing asymmetry

Arm-swing is thought to optimize dynamic stability and minimize the energy expenditure of gait (4-6, 8, 48). Coupling-effects between the upper-limbs and their contralateral lower-limb remaining from our evolutionary quadrupedal past exemplify further the importance of appropriate arm-swing for locomotion (49). Previous studies investigating arm-swing asymmetry in healthy cohorts posited this asymmetry to be linked to handedness, where a right-hand dominant population were thought to reduce their preferred side for activities requiring fine-motor skills (15, 16). No association however was found between handedness and arm-swing asymmetry. An explanation from this for the presence of left-side dominant arm-swing in left-handed individuals was cultural mediation via a right-hand dominant society, in what was propositioned with relation to the ‘Gunslinger gait’ (15, 50). This would however theoretically leave left-handed individuals at a distinct disadvantage in terms of gait stability and energy efficiency if it were the case (6, 29).

The findings of the current study suggest that arm-swing asymmetry works as a counterbalance to asymmetry in lower-limb mechanical work. Aim 1 findings indicate a right-side predominance for negative mechanical work compared to left-side double-support. Soo & Donelan (2012) found the coordination of push-off and collision work determined the magnitude of mechanical work conducted during step-to-step transitions, exemplifying the necessity for equal coordination between limbs. A source for this uncoordinated push-off and collision work during step-to-step transitions was identified in the current study as sidedness which was reduced in the presence of higher N_k_. O’Reilly & Federolf (2020) revealed that the magnitude of left – right side weight transfer significantly moderated the relationship between upper- and lower-limb postural corrections also, thus exemplifying the potential role of sidedness. Asymmetries in joint moment contributions as high as 10% have been reported in healthy adults (51), explaining perhaps this increased need for angular momentum cancellation in some healthy individuals.

Aim 2 findings support this suggestion further by demonstrating that with a greater N_k_ value, dASI and W_ss_became increasingly more asymmetrical. A higher N_k_ is representative of participants who found very slow walking more challenging and by virtue of the aim 1 findings we have sufficient insight to determine that sidedness played a role in this increased difficulty. The aim 2 findings go further by demonstrating that cross-symmetries between the upper- and lower-limbs are actively compensated for by the neuromuscular system during a challenging motor task. In hemiparetic gait, unaffected side arm-swing is typically dominant and acts as a counterbalance to increased angular momentum in the affected lower-limb (12). The cross-over interaction demonstrated in the current study illustrates that this compensation strategy was utilised to a much smaller extent by some participants as right-side dominant arm-swing had an opposing effect to that of left-side dominant arm-swing. From this, it can be determined that arm-swing asymmetry is positively related to the degree of asymmetry present in lower-limb mechanical work. One suggestion for this asymmetric control strategy is the presence of cerebral lateralization of lower-limb motor control (i.e. footedness). Footedness has been noted to differ in terms of its cerebral lateralization to that of handedness with most right- and left-handed individuals in fact demonstrating right footedness (52, 53). This may explain how most studies have found that healthy, left-handed individuals exhibited a left-side dominant arm-swing (10, 15, 16). Interestingly, no association between spatiotemporal gait parameters and arm-swing asymmetry was found in previous research (16). Moreover no differences between dominant and non-dominant leg symmetry in terms of temporal and kinematic data have been found in the literature (54), however differences in terms of EMG profiles (55) and overall positive work (56) are cited thus favouring this proposition. Formal evidence for this relationship is required and future studies should look to confirm this association.

### 2.2 Open- and closed-loop control of whole-body angular momentum

The observations made by Robert et al. (2009) concerning the differential control of the WBAM during the single- and double-support phase of gait was supported by the findings of this study. N_k_ had a significant negative effect during double-support illustrating an adjustive role while although insignificant, a contrasting positive effect during single-support was found supporting the stabilizing role proposed. Wu et al. (2019) found a high variability in stance time at very slow walking speeds in healthy adults accompanied by relatively consistent swing-times, also supporting this differential control between phases. How the balance control mechanisms incorporated in WBAM regulation differ across walking-speeds has not been investigated thoroughly. Interdependencies between such gait balance control strategies relevant to step-to-step transitions (i.e. push-off modulation, lateral ankle and foot placement mechanisms) have been identified in the literature (17, 57). The interdependencies between lateral ankle and foot placement mechanisms were found to be consistent with varied perturbation direction, however only a weak association with the push-off mechanism was found. They suggested that the push-off mechanism may in fact have an indirect effect through modulation of other gait parameters. In O’Reilly & Federolf (2020), the effect of the baseline push-off mechanism on gait coordination during a trial of perturbation varied with walking-speed in a manner that was subject-specific. This finding was supported by previous research that found double-support phase WBAM regulation to vary with walking-speed but single-support mechanisms to be speed invariant (58). Due to its sensitivity to stance/swing-time modulation, stride-time variability is a potential parameter that may fit the suggestion made in (57) which is said to have a quadratic relationship with walking-speed (59). The results of the current study add to these discussed findings further by presenting a protocol for investigating this indirect effect on such gait parameters while also suggesting that sidedness should be considered when evaluating dynamic balance control. Further research is required to establish the relationship between the various balance control mechanisms across walking-speeds and in the context of sidedness.

## 3. Conclusion

Separate control mechanisms were revealed through an investigation into the association between the neuromuscular control of muscle synergies and step-to-step mechanical work at the start and end points of single- and double-support phases. Healthy adults walking at a very slow speed resorted to greater ipsilateral collision work as a compensation strategy for the asymmetrical work contributions on both sides during both phases of gait. A functional role for arm-swing asymmetry was presented in counterbalancing lower-limb asymmetries that is thought to be related to footedness. Future studies should look to confirm the association between footedness and arm-swing asymmetry and elucidate the association between balance control mechanisms across walking-speeds and the effect of sidedness on these mechanisms.

## 4. Conflicts of interest

The author declares no known conflicts of interest were present in this study.

## Notes

### Competing Interest Statement

The authors have declared no competing interest.

